# Characterizations of SARS-CoV-2 mutational profile, spike protein stability and viral transmission

**DOI:** 10.1101/2020.05.03.066266

**Authors:** Sayantan Laha, Joyeeta Chakraborty, Shantanab Das, Soumen Kanti Manna, Sampa Biswas, Raghunath Chatterjee

## Abstract

The recent pandemic of SARS-CoV-2 infection has affected more than 3.0 million people worldwide with more than 200 thousand reported deaths. The SARS-CoV-2 genome has a capability of gaining rapid mutations as the virus spreads. Whole genome sequencing data offers a wide range of opportunities to study the mutation dynamics. The advantage of increasing amount of whole genome sequence data of SARS-CoV-2 intrigued us to explore the mutation profile across the genome, to check the genome diversity and to investigate the implications of those mutations in protein stability and viral transmission. Four proteins, surface glycoprotein, nucleocapsid, ORF1ab and ORF8 showed frequent mutations, while envelop, membrane, ORF6 and ORF7a proteins showed conservation in terms of amino acid substitutions. Some of the mutations across different proteins showed co-occurrence, suggesting their functional cooperation in stability, transmission and adaptability. Combined analysis with the frequently mutated residues identified 20 viral variants, among which 12 specific combinations comprised more than 97% of the isolates considered for the analysis. Analysis of protein structure stability of surface glycoprotein mutants indicated viability of specific variants and are more prone to be temporally and spatially distributed across the globe. Similar empirical analysis of other proteins indicated existence of important functional implications of several variants. Analysis of co-occurred mutants indicated their structural and/or functional interaction among different SARS-COV-2 proteins. Identification of frequently mutated variants among COVID-19 patients might be useful for better clinical management, contact tracing and containment of the disease.

## INTRODUCTION

In December 2019, Chinese government reported several human pneumonia cases in Wuhan city and designated the disease as coronavirus disease 2019 (COVID 19) [1]. Major symptoms of COVID-19 include fever, cough, dyspnoea and muscular soreness. There were some patients with COVID 19, where atypical symptoms like diarrhoea and vomiting were also found [2–4]. Whole genome sequencing showed that the causative agent is a novel coronavirus and initially termed as 2019-nCoV [4, 5]. Later on, International Committee on Taxonomy of Viruses (ICTV) officially designated the virus as SARS-CoV-2. WHO on March 11, 2020 has declared COVID-19 outbreak as a global pandemic [6].

Corona-viruses are class of genetically diverse viruses found in a wide range of host species like mammals and birds [7, 8]. SARS-CoV-2 is an enveloped virus and comprises a positive sense single strand RNA genome of ∼30 kb [9]. This SARS-CoV-2 also belongs to the genus *betacoronavirus* like SARS-CoV and MERS-CoV. Primarily, it was thought to cause infections in birds and other mammals but recent outbreaks have clearly revealed the ability of coronaviruses to cross species barriers and human transmission [10]. Coronaviruses carry the largest genome among all RNA viruses and each viral transcript consists of a 5’-cap structure and a 3’ poly-A tail [11]. After entry to the host cell, the genomic RNA is translated to produce non-structural proteins (nsps) from two open reading frames (ORFs). On the other hand, the viral genome is also used as template for replication and transcription via RNA-dependent RNA polymerase activity. In the intermediate stage, negative strand RNA intermediates are produced to serve as template for positive sense genomic RNA and sub-genomic RNA synthesis. These shorter sub-genomic RNAs encode the structural proteins i.e. Spike, Envelope, Membrane and Nucleocapsid protein and several other accessory proteins [12–14].

The mutation rate for RNA viruses is drastically high and this higher mutation rate is correlated with the virulence which is beneficial for viral adaptation [15]. The SARS-CoV-2 genome has a capability of gaining rapid mutations as the virus spreads [16]. The advantage of increasing amount of whole genome sequence data of SARS-CoV-2 intrigued us to explore the mutation profile across the genome, to check the genome diversity and to investigate the consequences of those mutations in stability and transmission.

In this present study, we used ∼ 660 complete SARS-CoV-2 genome data from NCBI virus database (16^th^ April, 2020) for in-silico analysis. We performed gene and protein sequence alignment, and characterized the mutation status of all genes. The most conserved and variable regions were recognized for all genes along with synonymous and non-synonymous changes. As non-synonymous changes dictate the altered amino acid composition, a collection of all mutations for each protein has been determined. We catalogued these substitutions for all proteins and identified different variants that are prevalent in nature. We also evaluated the impact of mutating spike glycoprotein in protein stability, viral transmission, adaptability and diversification. This brief characterization of SARS-CoV-2 variants and functional impact analysis of those variants could lead to better clinical management of the COVID-19 pandemic.

## MATERIALS AND METHODS

### Collection of SARS-CoV-2 genomic sequences

Total 664 SARS-CoV-2 whole genome sequences were downloaded from NCBI Virus repository (https://www.ncbi.nlm.nih.gov/labs/virus) as of April 16, 2020. Partially sequenced genomes and sequences with ambiguous sites were filtered out for further analysis. Number of sequences from each country is as follows: Australia:1, Brazil: 1, China: 57, Colombia: 1, France: 1, Greece: 4, India: 2, Iran: 1, Israel: 2, Italy: 2, Nepal: 1, Pakistan: 2, Peru: 1, South Africa: 5, Spain: 11, Sweden: 1, Taiwan: 3, Turkey: 1, USA: 565 and Vietnam: 2.

### Identification of variable sites

We have aligned nucleotide and amino acid sequences of ORF1ab, ORF3a, ORF6, ORF7a, ORF8, ORF10, envelop (E), membrane (M), nucleocapsid (N) and surface glycoprotein (S) using MUSCLE multiple sequence alignment algorithm in MEGA-X [17]. We tabulated the number of variable, singleton, parsimony informative sites at both gene and protein levels. After removing the ambiguous and deleted residues, we determined the amino acid substitutions in all ten proteins of SARS-CoV-2 genome. Frequently mutated residues are those that showed mutation in 1.5% at least (10 strains) of the strains. Co-occurring mutations are determined considering all frequently mutated residues for S, N, ORF3a, ORF8 and ORF1ab proteins.

Since it has been well established that bats as well as pangolins may be the sources of the original transmission of the virus in humans [18, 19], we were interested to note the amino acid residues in the key mutational sites in Bat and pangolin coronavirus protein sequences. The protein sequences of Bat coronavirus RaTG13 (GenBank Accession ID MN996532.1), Pangolin coronavirus (MT072864.1) and two SARS-CoV strains TW11 (AY502924.1) and GD01 (AY278489.2) were downloaded from NCBI (https://www.ncbi.nlm.nih.gov/protein). These were aligned with the analogous protein sequences of SARS-Cov2 in MEGA-X, and the residues at the frequently mutated sites of the respective viral proteins were observed.

### Protein structural analyses

Cryo-EM three-dimensional structures of SARS-CoV-2 spike glycoprotein have been recently made available in RCSB Protein Data Bank (https://www.rcsb.org/) [20]. Three pdb structures i.e. 6VXX (Closed state) [21], 6VYB (Open state) [21] and 6VSB (Prefusion) [22] were downloaded. We used FoldX BuildModel function to construct the mutant 3D protein structures [23]. The differences in total energy, electrostatics, solvation energy etc. were calculated for all the closed, open state as well as prefusion spike protein mutant 3D models. All three cryo-EM structures have some missing residues in the available PDBs, and are not considered for stability calculation or further structural analysis. Structural analyses and figures have been generated in Discovery studio 2020 (Dassault System BIOVIA Corp) [24].

### Phylogeny, and Spatial and temporal distribution of SARS-CoV-2 strains

To study the impact of non-synonymous mutations in the transmission and viability of the newly emerging SARS-CoV-2 sequences across the world, we looked into Nextstrain tool, a powerful visualization tool comprising >10,000 SARS-CoV-2 samples to study the evolution of various pathogens (https://nextstrain.org/) [25]. The frequency of occurrence, the date of collection and corresponding geographical location of the mutant strains were noted on 27^th^ April, 2020. Please note that the genotype information on Nextstrain are not consistent on different dates, at least for some residues. Major amino acid substitutions were mapped and visualized in the phylogeny from Nextstrain Next hCoV-19 App. Minor (less frequently) mutated residues that showed variations in Nextstrain data compared to NCBI virus database are excluded from the analysis. Amino acid residues of ORF1ab were split as ORF1a from 1-4400 and the rest as ORF1b in Nextstrain Next hCoV-19 App.

### Empirical analysis of functional implication of mutations

To understand the implications of the amino acid substitutions in the mutants, charge state (at neutral pH), Kyte-Doolittle hydropathy indices [26] and Grantham's mean chemical difference indices, which takes into account side-chain chemical structure, volume and polarity [27], were compared. Stabilities of their side-chains between exposed and buried forms were compared using apparent partition energies as reported by Wertz and Scheraga [28]. The typical contributions of a putative H-bond and salt-bridge towards protein stabilities were assumed to be in the range of 0.5-1.5 kcal/mole [29–31] and 3 kcal/mole [32] based on previous studies on protein folding.

## RESULTS

### Synonymous and non-synonymous mutations

After aligning the individual ORFs of SARS-CoV-2 genome, we identified mutations both at gene and protein levels (Table 1). At the gene level, the number of mutations per 100 bases were found to be relatively high in N, ORF10, ORF6, ORF7a, ORF8 and ORF3a, suggesting that these genes may be more prone to mutations as compared to others. Among the structural proteins, M and E proteins contained the least variability, which could indicate that these proteins may be associated with housekeeping functions and consequently have a greater resistance to mutations. Looking into the changes per 100 amino acids for each of the proteins (Table 1), we observed that ORF3a exhibited the highest mutability, closely followed by N and ORF8. While looking into the synonymous and non-synonymous changes, we have found that the ORF1ab and spike protein contained the largest number of non-synonymous mutations (Table 1). When we normalized w.r.t to the length, ORF3a, ORF8 and N exhibited relatively high number of non-synonymous changes.

**Table 1:** Nucleotide and protein alignment of SARS-CoV-2 genes

| Name | Total No of Samples | No. of nucleotides | Variable sites | Variable Sites per 100 bp | Synonymous | Non Synonymous | Singleton Informative Site per 100 bp | Parsimony Informative Site per 100 bp | No. of Amino acids | Variable Site | Variable Sites per 100 aa | Singleton Informative Site per 100 aa | Parsimony Informative Site per 100 aa |
| --- | --- | --- | --- | --- | --- | --- | --- | --- | --- | --- | --- | --- | --- |
| **Envelope Protein** | 664 | 228 | 4 | 1.75 | 2 | 2 | 1.32 | 0.44 | 76 | 2 | 0.88 | 0.88 | 0.00 |
| **Membrane Protein** | 649 | 669 | 7 | 1.20 | 2 | 5 | 0.60 | 0.60 | 223 | 5 | 0.75 | 0.45 | 0.30 |
| **Nucleocapsid Protein** | 664 | 1260 | 50 | 3.97 | 24 | 26 | 2.54 | 1.43 | 420 | 26 | 2.06 | 1.43 | 0.63 |
| **ORF10** | 660 | 117 | 4 | 3.42 | 2 | 2 | 2.56 | 0.85 | 39 | 2 | 1.71 | 1.71 | 0.00 |
| **ORF1ab** | 639 | 21291 | 383 | 1.80 | 148 | 235 | 1.34 | 0.46 | 7097 | 235 | 1.10 | 0.83 | 0.28 |
| **ORF3a** | 660 | 828 | 28 | 3.38 | 8 | 20 | 2.05 | 1.33 | 276 | 20 | 2.42 | 1.33 | 1.09 |
| **ORF6** | 657 | 186 | 6 | 3.23 | 3 | 3 | 1.61 | 1.61 | 62 | 3 | 1.61 | 1.08 | 0.54 |
| **ORF7a** | 658 | 366 | 9 | 2.73 | 3 | 6 | 1.91 | 0.82 | 122 | 6 | 1.64 | 1.09 | 0.55 |
| **ORF8** | 661 | 366 | 9 | 2.46 | 1 | 8 | 1.37 | 1.09 | 122 | 8 | 2.19 | 1.09 | 1.09 |
| **Spike Protein** | 643 | 3822 | 74 | 1.96 | 30 | 44 | 1.52 | 0.44 | 1274 | 44 | 1.13 | 0.92 | 0.21 |

### Frequently mutated amino acids in SARS-CoV-2 proteins

Considering the altered protein sequences due to non-synonymous changes, we next focussed on amino acid substitutions among all proteins of the SARS-CoV-2 genome (Supplementary Table 1). An abridged version of the table containing only those substitutions that have been observed in a minimum of 10 isolates (∼1.5% of the total isolates) is provided in Figure 1. Among the structural proteins, E and M showed most conserved structures across all the viral genomes under consideration, with substitutions at two sites of each E and M proteins only in 1 and 5 isolates respectively. Upon examining the S proteins, we find several mutations; nevertheless most of these substitutions are perceived only in a single isolate, with notable exceptions being the D614G. There have been 264 (41%) instances of D614G, suggesting its pivotal role in regards to the protein stability and other key characteristics. Among the other changes, V483A, L5F, Q675H, H655Y and S939F occurred in 6, 5, 3, 2 and 2 isolates respectively. The N protein also depicted substitutions R203K and G204R.

**Figure 1.**
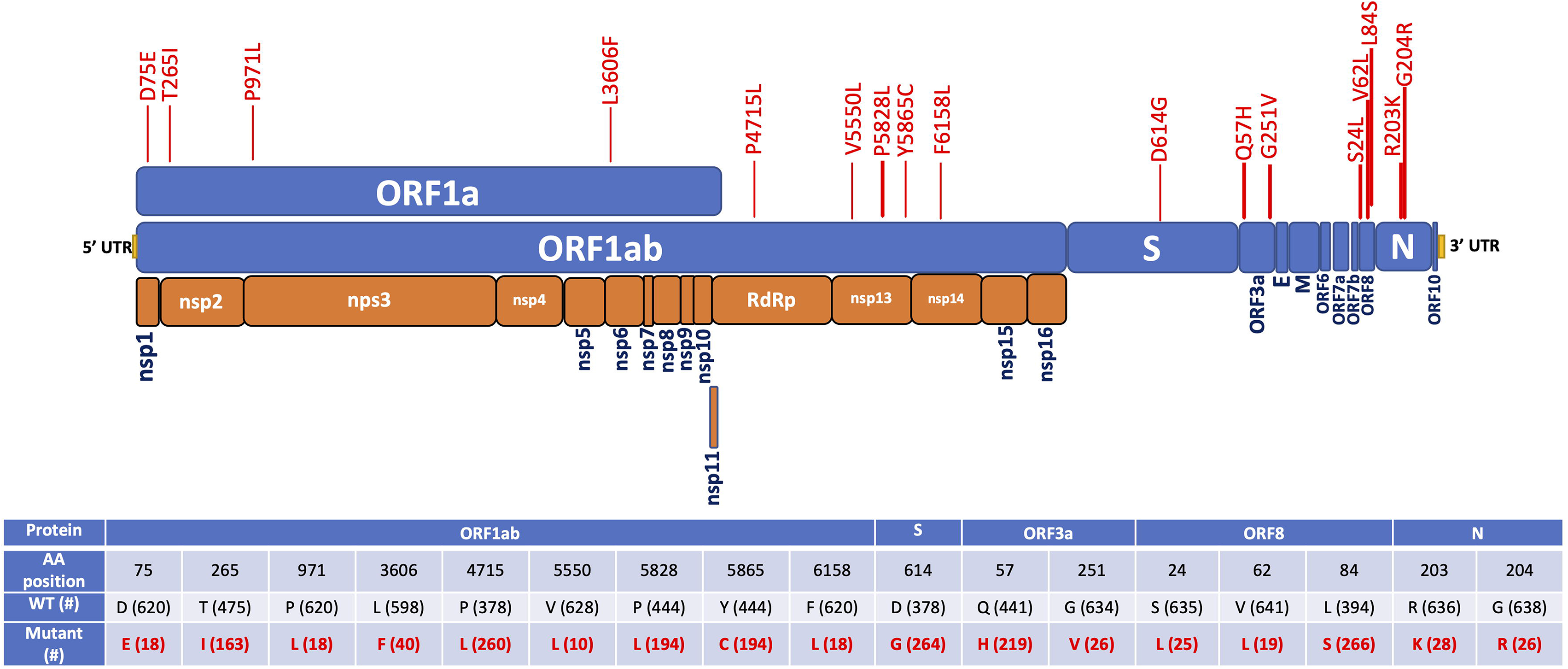
Frequently mutated residues are plotted on the respective proteins of SARS-CoV-2. Tabular presentation depicted the number of occurrences (#) of wildtype and mutant residues in ∼660 samples.

Among the non-structural proteins, ORFs 6, 7a and 10 share similar behaviour to E and M proteins with them being mostly conserved. In contrast, ORF3a exhibits non-synonymous mutations, the majority of which have mostly been distributed at 2 residues (Q57H and G251V). Mutations in ORF8 showed a major substitution at L84S and an accompanied change of V62L. Another substitution, S24L is observed in 25 samples out of the 660 sequences analysed. We move on to the largest encoded SARS-CoV-2 protein, ORF1ab that encodes replicase polyproteins required for viral RNA replication and transcription [33]. ORF1a and ORF1b encode two polypeptides, pp1a and pp1ab, and finally processed into 16 nsps (Figure 1) [33, 34]. Majority of the non-synonymous mutations at ORF1ab leads to amino acid changes at the 265^th^, 4725^th^, 5828^th^ and 5865^th^ residues (T265I, P4715L, P5828L and Y5865C). Notably, L5828P and C5865Y occurred simultaneously in all strains, suggesting possible functional relationship between these two residues.

### Identification of prevalent SARS-CoV-2 variants

To identify the variants that are prevalent during the course of time, we next determined the frequently mutated residues that occurred at least in 1.5% of the samples. The combined analysis of all proteins with these frequently mutated residues identified 20 possible SARS-CoV-2 variants, among which 15 variants comprised more than 97% of the analysed sequences having frequency >1% (Table 2). Apart from the wild type variant (13.3%), other frequent SARS-CoV-2 variants are ORF8:L84S / ORF1ab:P5828L/Y5865C (30.6%), S:D614G / ORF3a:Q57H / ORF1ab:T265I/P4715L (20.4%), S:D614G / ORF3a:Q57H / ORF1ab:P4715L (7.2%), ORF8:L84S (4.6%) and S:D614G / ORF1ab:P4715L (4.6%). We noted that G251V of ORF3a, V62L of ORF8, D75E, P971L, P5828L, Y5865C and F6158L of ORF1ab substitutions occurred only with the D614 wild type variant of the S protein. While, substitutions R203K and G204R of N protein, Q57H of ORF3a, S24L of ORF8, T265I, P4715L and V5550L of ORF1ab occurred only with the D614G mutant of S protein. Co-occurrences of these mutations might have implications in direct structural interactions or indirect regulations of these proteins on survivability of the virus. To further validate these finding, we visualized these major substitutions by observing the phylogeny of SARS-CoV-2 in Nextstrain, which contains a curated database of more than 10,000 SARS-CoV-2 sequences, depicted in the form of phylogenetic trees (Figure 2).

**Figure 2.**
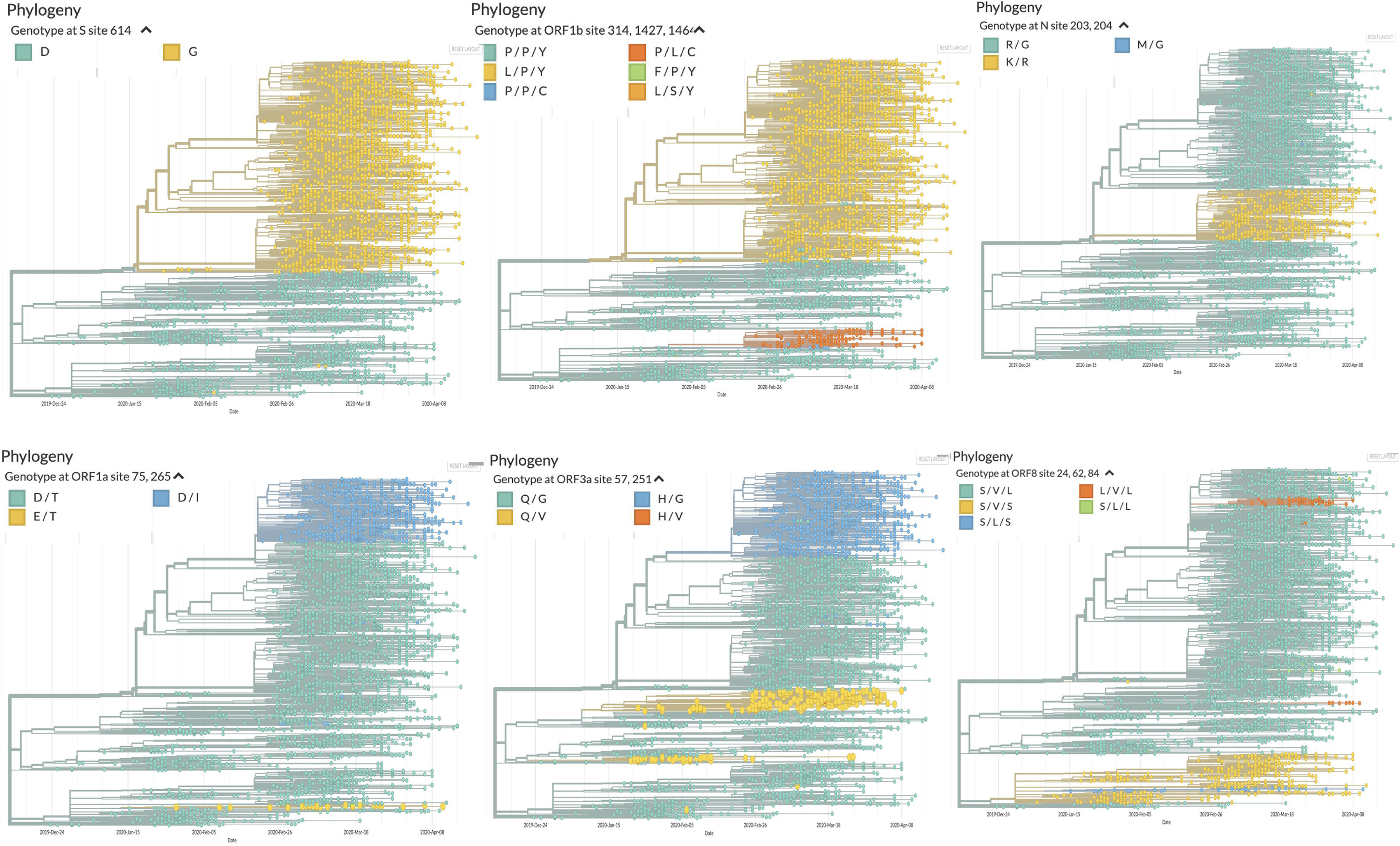
Phylogeny (generated from Nextstrain) with colour coding for the genotypes at **A.** 614^th^ amino acid position of Spike glycoprotein (S), **B.** 314, 1427 and 1464^th^ positions of ORF1b, **C.** 203 and 204^th^ positions of Nucleocapsid (N), **D.** 75 and 265^th^ of ORF1a, **E.** 57 and 251^st^ positions of ORF3a and **F.** 24, 62 and 84^th^ positions of ORF8 proteins.

**Table 2:** Strains with co-occurring major mutations in N, S, ORF3a, ORF8 and ORF1ab proteins

| Proteins (Positions) | N(203,204) |  | S(614) | orf3a(57,251) |  | orf8(24,62,84) |  |  | orf1ab (75,265,971,3606,4715,5550,5828,5865,6158) |  |  |  |  |  |  |  |  | # of isolates | Freq. (%) |
| --- | --- | --- | --- | --- | --- | --- | --- | --- | --- | --- | --- | --- | --- | --- | --- | --- | --- | --- | --- |
| Amino acid position | 203 | 204 | 614 | 57 | 251 | 24 | 62 | 84 | 75 | 265 | 971 | 3606 | 4715 | 5550 | 5828 | 5865 | 6158 |  |  |
| SARS-CoV-2 | R | G | D | Q | G | S | V | S | D | T | P | L | P | V | L | C | F | 186 | 30.6 |
|  | R | G | G | H | G | S | V | L | D | I | P | L | L | V | P | Y | F | 124 | 20.4 |
|  | R | G | D | Q | G | S | V | L | D | T | P | L | P | V | P | Y | F | 81 | 13.3 |
|  | R | G | G | H | G | S | V | L | D | T | P | L | L | V | P | Y | F | 44 | 7.2 |
|  | R | G | D | Q | G | S | V | S | D | T | P | L | P | V | P | Y | F | 28 | 4.6 |
|  | R | G | G | Q | G | S | V | L | D | T | P | L | L | V | P | Y | F | 28 | 4.6 |
|  | K | R | G | Q | G | S | V | L | D | T | P | L | L | V | P | Y | F | 23 | 3.8 |
|  | R | G | D | Q | V | S | V | L | D | T | P | F | P | V | P | Y | F | 20 | 3.3 |
|  | R | G | G | H | G | L | V | L | D | I | P | L | L | V | P | Y | F | 20 | 3.3 |
|  | R | G | D | Q | G | S | L | S | E | T | L | L | P | V | P | Y | L | 17 | 2.8 |
|  | R | G | D | Q | G | S | V | L | D | T | P | F | P | V | P | Y | F | 11 | 1.8 |
|  | R | G | G | H | G | S | V | L | D | I | P | L | L | L | P | Y | F | 8 | 1.3 |
|  | R | G | D | Q | V | S | V | L | D | T | P | L | P | V | P | Y | F | 5 | 0.8 |
|  | R | G | D | Q | G | S | V | S | D | T | P | F | P | V | L | C | F | 4 | 0.7 |
|  | R | G | D | Q | G | S | V | S | D | T | P | F | P | V | P | Y | F | 3 | 0.5 |
|  | R | G | D | Q | G | S | L | S | E | T | L | L | P | V | P | Y | F | 1 | 0.2 |
|  | R | G | D | Q | G | S | L | S | D | T | P | L | P | V | P | Y | F | 1 | 0.2 |
|  | R | G | G | H | G | S | V | L | D | I | P | F | L | V | P | Y | F | 1 | 0.2 |
|  | K | G | G | H | G | S | V | L | D | I | P | L | L | V | P | Y | F | 1 | 0.2 |
|  | R | G | G | H | G | S | V | S | D | I | P | L | L | V | P | Y | F | 1 | 0.2 |
| Bat coronavirus RaTG13 | R | G | D | Q | G | S | V | S | D | T | P | V | P | V | P | Y | F |  |  |
| Pangolin coronavirus | R | G | D | Q | G | S | V | S | D | N | Q | V | P | V | P | Y | F |  |  |
| SARS coronavirus TW11 | R | G | D | Q | G | E | F | Y | D | T | V | V | P | V | P | Y | F |  |  |
| SARS coronavirus GD01 | R | G | D | Q | G | E | F | Y | D | T | V | V | P | V | P | Y | F |  |  |

\* Mutated residues are presented in red

We observed the variants that contained these specific substitutions are mostly clustered together. For S protein, proportion of the samples with D614G substitutions was roughly equal to that of wild type variant, which shows the adaptability of this substitution. For ORF1b, we assessed the substitutions at positions 314, 1427 and 1464, which corresponded to 4715^th^, 5828^th^ and 5865^th^ residues of ORF1ab. Mapping of all frequently mutated residues of ORF1a and ORF1b on the phylogeny of SARS-CoV-2 is presented in Supplementary Figure 1. As observed previously, substitutions at 5828^th^ and 5865^th^ position co-occurred even in this large sample set of Nextstrain data. Viral variants with residues L/P/Y, P/P/Y and P/L/C dominate the bulk of the sequences, which shows that the conjoined mutations (either P/Y or L/C) at 5828^th^ and 5865^th^ residues are linked with increasing survivability (Figure 2). Interestingly, we could not overlook the fact that the two representations of the mutations for both S protein and ORF1b were remarkably co-occurring. We see that those variants that have D614G substitutions in S also have L/P/Y residues at 314, 1427 and 1464 positions of ORF1b. It suggests that these residues in S and ORF1b, irrespective of whether they have occurred simultaneously, show good viability (Figure 2). When focusing on the D614 residue in S protein, the most prevalent variant are with P/P/Y at all the three positions of ORF1b, with exception of few variants with P/L/C residues. Looking into the mutation profile of N, we find that the wild type variants (R/G at 203 and 204 positions) appear to be predominant. Comparison with mutation profile of S protein identified co-occurrence of 614G variant with K/R at 203^rd^ and 204^th^ positions of N protein (Figure 2). We checked the mutation status at positions 75 and 265 of ORF1a protein. We withheld the inclusion of the 971^st^ site here, as we have seen that substitutions at this site and the 75^th^ residue are co-linked, i.e. they are mostly identical (Figure 2). We notice that wild type variants form the majority with few isolates of D75E mutants. Fairly good number of samples with T265I substitutions of ORF1a are also observed having D614G substitutions of S protein. Comparing the mutational profile with the ORF3a at 57^th^ and 251^st^ positions, we again find a stark resemblance to both these profiles. We see that the D/I variant in ORF1a mostly went hand-in-hand with H/G variants in ORF3a (Figure 2). Viewing the mutational profile of ORF8 with respect to positions V62L and L84S occurred mostly with the D614 of S protein, while S24L occurred with the G variants, also observed in our analysis with ∼660 samples. Overall, the mutation profile that we identified with 664 samples showed excellent concordance with the Nextstrain data comprising ∼10,000 samples.

We compared these frequently mutated residues with the corresponding protein sequences of Bat coronavirus RaTG13, Pangolin coronavirus, SARS coronavirus TW11 and GD01 (Table 2). All these frequently mutated residues completely matched with Bat coronavirus RaTG13. The notable exceptions in here being mismatches at positions 265, 971 and 3606 of ORF1a in case of pangolin coronavirus and mismatches in all three residues of ORF8 in both the SARS-CoV isolates.

### Empirical analysis of changes in structure and stability parameters

Parameters for 17 frequently mutated residues and V483A substitution at the receptor binding domain of S protein that may affect the protein structures are presented in Table 3. Only 3 out of these 18 substitutions were associated with any change in the charge of the side chain. The change in apparent side-chain partitioning energy varied from −2.79 to 3.13 kcal/mole. Maximum expected difference in number of H-bond and salt bridges associated with mutations were 4 and 1, respectively. While only 3 variants could have change in salt-bridge interactions, 8 of them could potentially have difference in H-bonding reflecting the fact that average energy associated with a salt-bridge interaction is much higher.

**Table 3:**
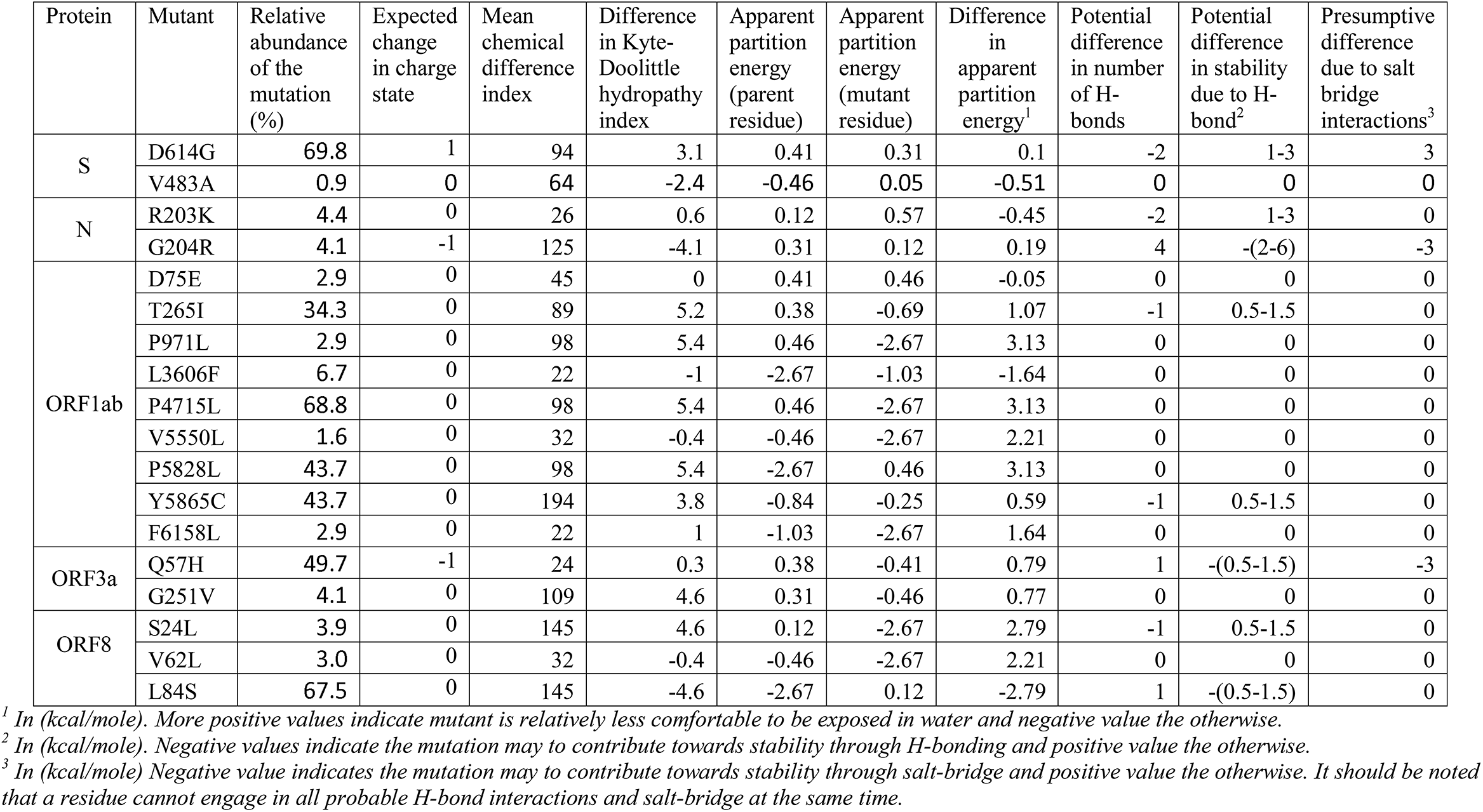
Potential implications of SARS-COV2 mutations on viral protein structure and stability.

The relative abundance of a mutation can be taken as surrogate of viral viability, which would be dictated by the effect of the mutation on protein stability and its function with respect to specific biomolecular events during host-pathogen interaction. The relative abundance of the D614G mutant (69.8%) was the highest. The partitioning energy difference was minimal (0.1 kcal/mole) indicating that unless the sidechain of D614 was involved in any consequential H-bond, this mutant could be as viable as the parent strain. The analysis of the larger Nextstrain dataset indicated that D614G mutation is significantly more prevalent, indeed. The high relative abundance (67.5%) of L84S mutant (ORF8) could be due to additional H-bond or favorable partitioning energy. T265I and Y5865C mutants of ORF1ab associated with removal of an alcoholic -OH group, which is often associated with modest contribution to protein stability [29–31] showed similar relative abundance of 34.3% and 43.7%, respectively. However, S24L mutant (ORF8) showed significant decrease in abundance. This may be attributed to more unfavorable change in apparent partitioning energy (by ∼2 kcal/mole) due to significant difference in chain length of serine and leucine. The significant decrease in abundance of G204R (N protein) in spite of potential for additional H-bonding may also be attributed to significantly larger chain length. However, the V483A (S protein), V62L (ORF8), V5550L and D75E (ORF1ab) mutants with relatively low values of differences in Grantham's index or Kyte-Doolittle index and comparable H-bonding or salt-bridge capacity showed dramatic decrease in relative abundance. While P4715L and P5828L mutants showed relatively high abundance (39.2 and 29.2%, respectively), P971L showed only 2.7% abundance. Interestingly, both F6158L and L3606F with very low differences Grantham's index and reversal in the difference in apparent partitioning energy showed low abundance.

### Stability of mutant spike glycoproteins

We have encountered several different variants pertaining to S protein of SARS-CoV-2. Apart from D614G and some co-occurring mutations, other changes have been observed in a few cases, e.g., L5F occurring in 5 strains, V483A in 6 strains, while among others, most of these substitutions were observed in a single strain. By performing the stability analysis of spike glycoprotein for mutating residues that are available in all three pdb file, we found that some of the variants are stable in nature corresponding to negative total energies, calculated for both open, closed as well as prefusion models (Table 4, Supplementary Table 2). Among 22 analysed substitutions in S protein, 9 structures showed reduction in total free energy in all three conformations. Mutants S50L and H49Y showed the most reduction in total free energy, while all mutants with D614G substitutions showed stabilizing structure, suggesting its prevalent role in spike protein evolution. Interestingly, reduction in solvation polar energy was found in only 5 structures, including D614G mutant. Detailed information for the differences in energy for all residues are presented in Supplementary Table 2.

**Table 4:**
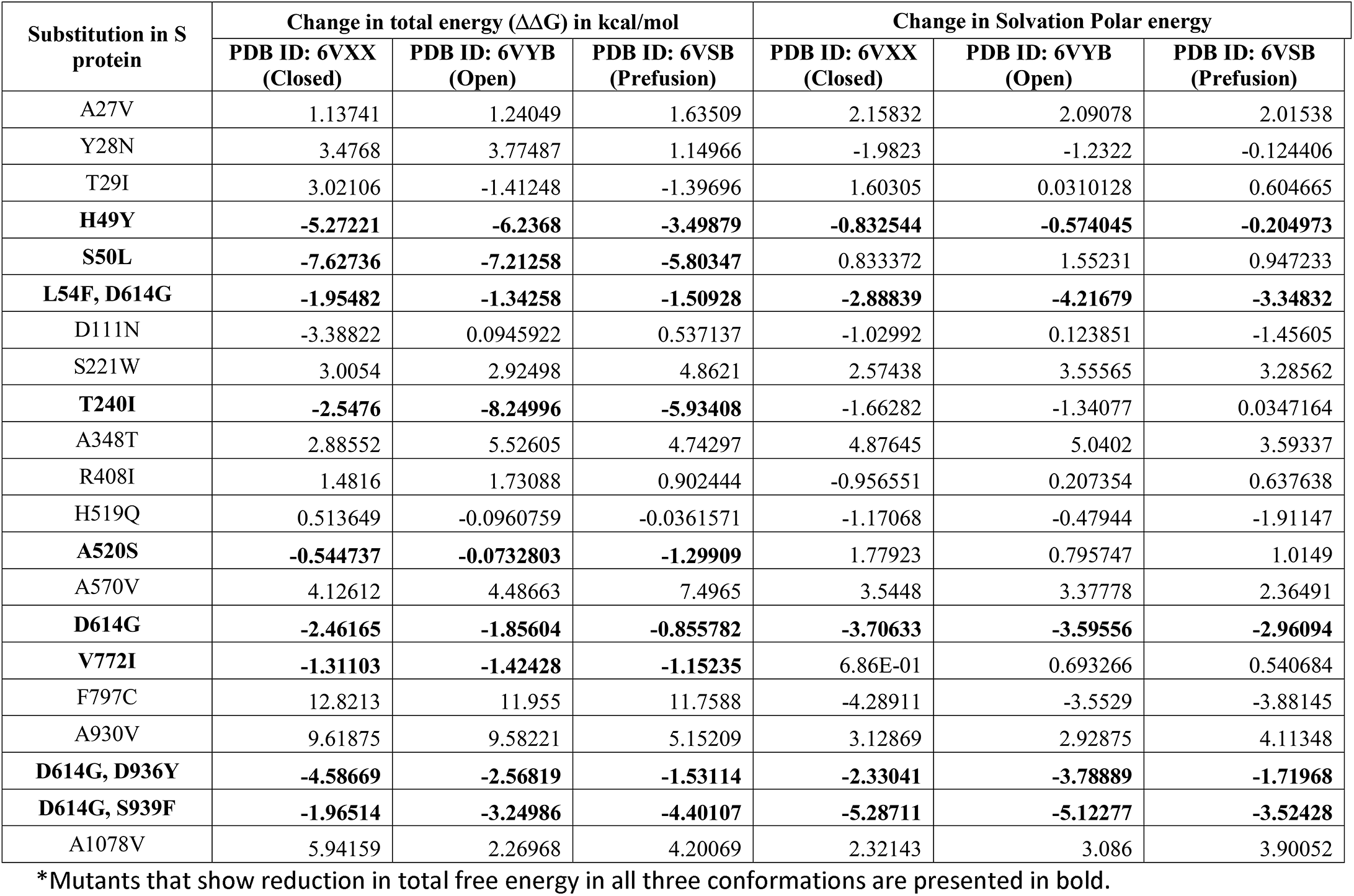
Total free energy and solvation polar energy changes of Spike glycoprotein mutants (Protein Stability analysis using FoldX)

| Substitution in S protein | Change in total energy ($\Delta\Delta G$) in kcal/mol | | | Change in Solvation Polar energy | | |
| --- | --- | --- | --- | --- | --- | --- |
|  | PDB ID: 6VXX (Closed) | PDB ID: 6VYB (Open) | PDB ID: 6VSB (Prefusion) | PDB ID: 6VXX (Closed) | PDB ID: 6VYB (Open) | PDB ID: 6VSB (Prefusion) |
| A27V | 1.13741 | 1.24049 | 1.63509 | 2.15832 | 2.09078 | 2.01538 |
| Y28N | 3.4768 | 3.77487 | 1.14966 | -1.9823 | -1.2322 | -0.124406 |
| T29I | 3.02106 | -1.41248 | -1.39696 | 1.60305 | 0.0310128 | 0.604665 |
| **H49Y** | **-5.27221** | **-6.2368** | **-3.49879** | **-0.832544** | **-0.574045** | **-0.204973** |
| **S50L** | **-7.62736** | **-7.21258** | **-5.80347** | 0.833372 | 1.55231 | 0.947233 |
| **L54F, D614G** | **-1.95482** | **-1.34258** | **-1.50928** | **-2.88839** | **-4.21679** | **-3.34832** |
| D111N | -3.38822 | 0.0945922 | 0.537137 | -1.02992 | 0.123851 | -1.45605 |
| S221W | 3.0054 | 2.92498 | 4.8621 | 2.57438 | 3.55565 | 3.28562 |
| **T240I** | **-2.5476** | **-8.24996** | **-5.93408** | -1.66282 | -1.34077 | 0.0347164 |
| A348T | 2.88552 | 5.52605 | 4.74297 | 4.87645 | 5.0402 | 3.59337 |
| R408I | 1.4816 | 1.73088 | 0.902444 | -0.956551 | 0.207354 | 0.637638 |
| H519Q | 0.513649 | -0.0960759 | -0.0361571 | -1.17068 | -0.47944 | -1.91147 |
| **A520S** | **-0.544737** | **-0.0732803** | **-1.29909** | 1.77923 | 0.795747 | 1.0149 |
| A570V | 4.12612 | 4.48663 | 7.4965 | 3.5448 | 3.37778 | 2.36491 |
| **D614G** | **-2.46165** | **-1.85604** | **-0.855782** | **-3.70633** | **-3.59556** | **-2.96094** |
| **V772I** | **-1.31103** | **-1.42428** | **-1.15235** | 6.86E-01 | 0.693266 | 0.540684 |
| F797C | 12.8213 | 11.955 | 11.7588 | -4.28911 | -3.5529 | -3.88145 |
| A930V | 9.61875 | 9.58221 | 5.15209 | 3.12869 | 2.92875 | 4.11348 |
| **D614G, D936Y** | **-4.58669** | **-2.56819** | **-1.53114** | **-2.33041** | **-3.78889** | **-1.71968** |
| **D614G, S939F** | **-1.96514** | **-3.24986** | **-4.40107** | **-5.28711** | **-5.12277** | **-3.52428** |
| A1078V | 5.94159 | 2.26968 | 4.20069 | 2.32143 | 3.086 | 3.90052 |

\*Mutants that show reduction in total free energy in all three conformations are presented in bold.

### Structural analysis of wild type and mutant spike glycoproteins

To further understand the implication of these mutants, we have analysed all three structures of S [21, 22, 35]. The spike glycoprotein is a homo-trimeric protein [21, 22, 35] having two subunits S1 and S2 in each monomer protruding from the viral surface. S1 subunit forms a budding head responsible for host–receptor binding, while S2 is mainly a stalk like structure that helps in fusion of viral and host membranes (Figure 3A). S proteins are cleaved at S1/S2 interface but remain non-covalently linked with each other in the prefusion state [35]. S1 subunit can further be divided into sub-domains namely N-terminal domain (NTD: residues 15-261), C-terminal domains 1, 2 and 3 (CTD1: residues 320-516; CTD2: residues 517-579; CTD3: residues 580-663) (Figure 3A).

**Figure 3.**
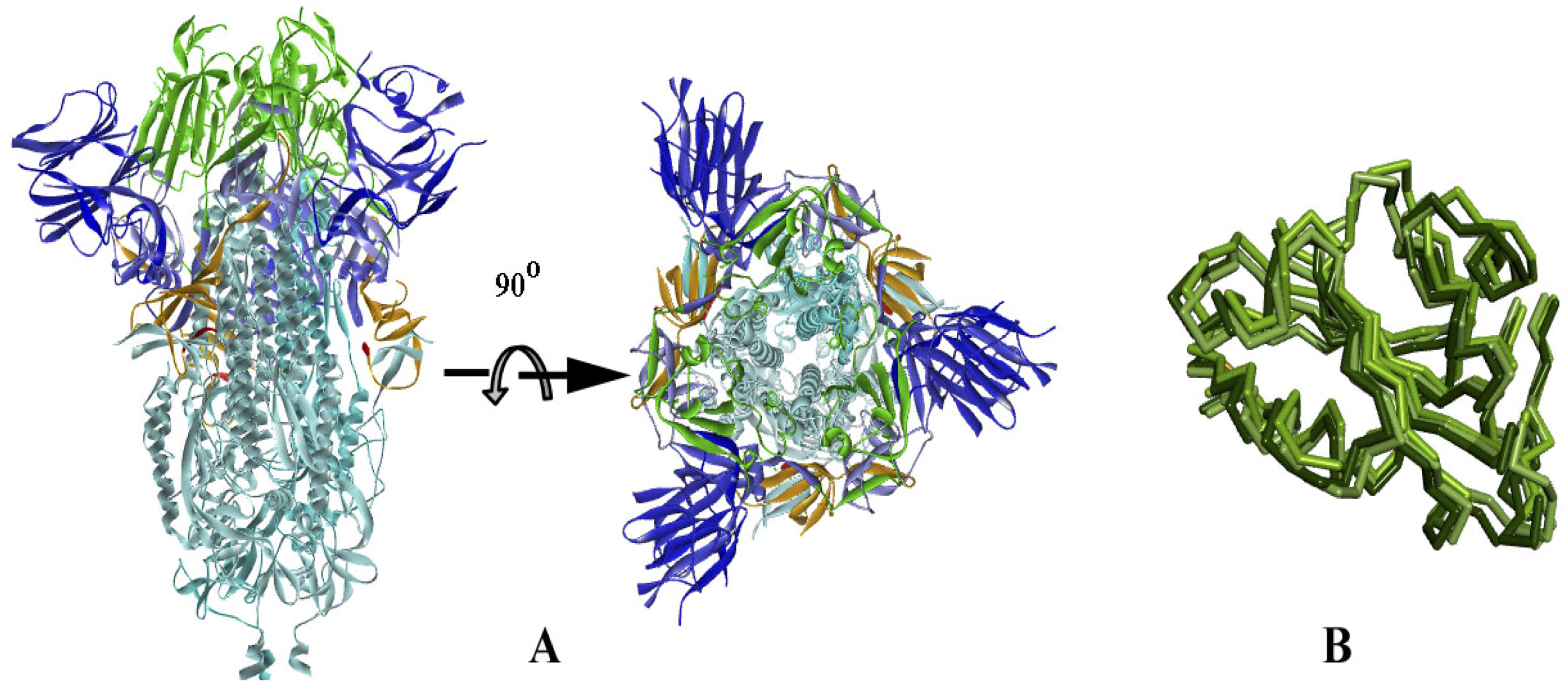
Ribbon diagram of Spike Protein (PDB_ID 6VXX), **A.** In two view; colour code: NTD blue, RBD green, CTD2 light blue, CTD3 orange, S1/S2 linker red and S2 sky blue. **B.** Superposition of RBD of DOWN/closed conformation (6VXX) with UP/open (6VYB) and per-fusion state (6VSB) in faded green colour ribbons.

CTD1, which is the main region of S protein for host-receptor interaction, is also termed as receptor binding domain (RBD) [21, 22, 35]. RBD undergoes conformational changes during receptor binding (human ACE2) that leads to a blossom of the S1 bud in an open or ‘UP’ conformation conducive for S-ACE2 interaction. Comparing the inert ‘DOWN’ and active ‘UP’ conformations (PDB_ID s: 6VXX and 6VYB respectively), it is found that RBD moves as a rigid-body in a hinge bending motion around its linker region with NTD and CTD2 with all atom rmsd for 198 residues is around 2.8 Å (Figure 3B). A similar change of conformation is also observed in prefusion state [22].

The miss-sense mutations in S protein are mainly single point mutations with few double mutations. All these mutations can be classified as stabilising and destabilising based on the free-energy changes (Table 4) of the in-silico generated mutant structures w.r.t the wild type variant [36] (Figure 1). Our study depicts that there are no stabilising mutations in the receptor binding domain RBD (Figure 4A, B). This observation indicates that the mechanism of S protein for a high affinity human ACE2 binding is unique in nature and any mutation (found till date) leads to an unstable structure and this could be correlated with lower viability of these mutation containing isolates.

**Figure 4.**
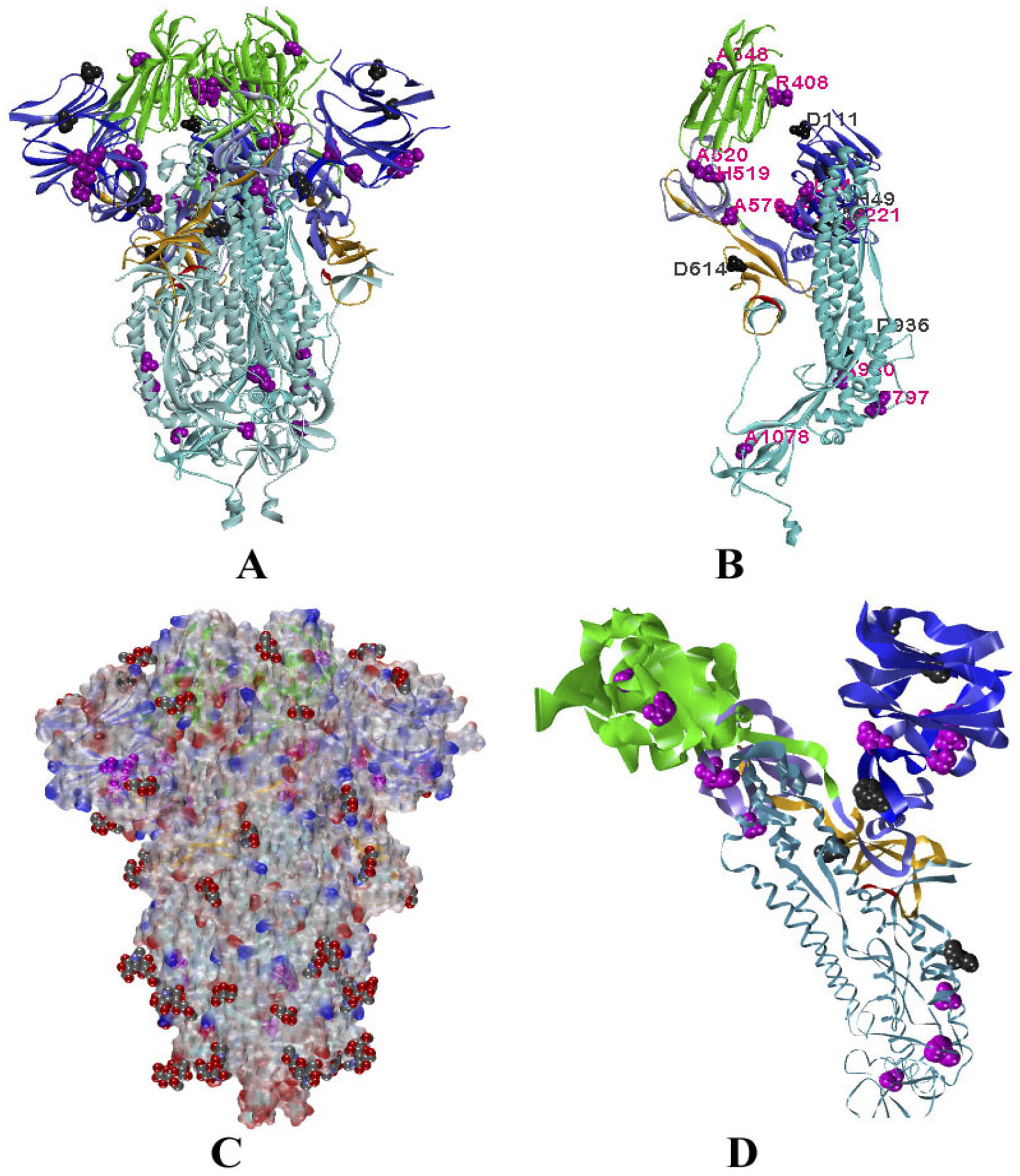
Location of mutations. **A.** Ribbon diagram of trimeric S protein with color code as described in Fig X1. The amino acids undergone to mutations are represented by Vander-wall radii with purple for destabilising/neutral mutation points and grey for stabilising ones. **B.** The monomeric unit of S-protein with label for the same amino acids as in A. **C.** A semi-transparent electro-static surface presentation of the S-protein with glycans as Van der waal presentation. **D.** Mutations in the monomeric unit of S protein, where ribbon size is proportional to average isotropic displacement of amino acid residue.

There are 42 miss-sense mutations found in S protein, we have considered 21 of them that are available in every monomer structure of S protein as well as in three pdbs. Out of these, 32 are in the S1 subunit and only 10 are found in S2 subunit. The cryo-EM structure of the protein (6VXX) shows high thermal parameter for the NTD and RBD (Figure 4D). High temperature factor of RBD could be correlated with its dynamic nature leading to the conformational switch between close and open states. It is also observed that most of the stabilizing mutations are in the NTD, which is an inherent unstable domain as depicted from high thermal parameter. This observation puts an open question whether virus adopts viability through mutations, stabilising the flexible NTD.

D614G substitution in CTD3 is found to be very stable and prevalent in nature. It occurs either as a single mutation or coupled with other mutations (L54F/D614G, D614G/D936Y and D614G/S939F). Surprisingly, L54F is a sort of neutral (in terms of free energy change) mutation, however, when coupled with D614G, the double mutant becomes a stable one (Table 4). The structural comparison of wild-type and in-silico generated D614G mutant shows that a change from Aspartic acid to Glycine alters electro-static potential of the surface of the protein (Figure 5A). This change creates a favourable environment in a hydrophobic pocket of the S protein (Figure 5B). Moreover, we have also observed that D614 is at the proximity of the hinge bending region (CTD2/NTD linker) of RBD (Figure 5C), therefore mutation of D to a small residue G without any side-chain might increase the flexibility for smooth switch over from inactive DOWN state to active UP state, makes the mutant containing variants more virulent in terms of its smoother binding with ACE2.

**Figure 5.**
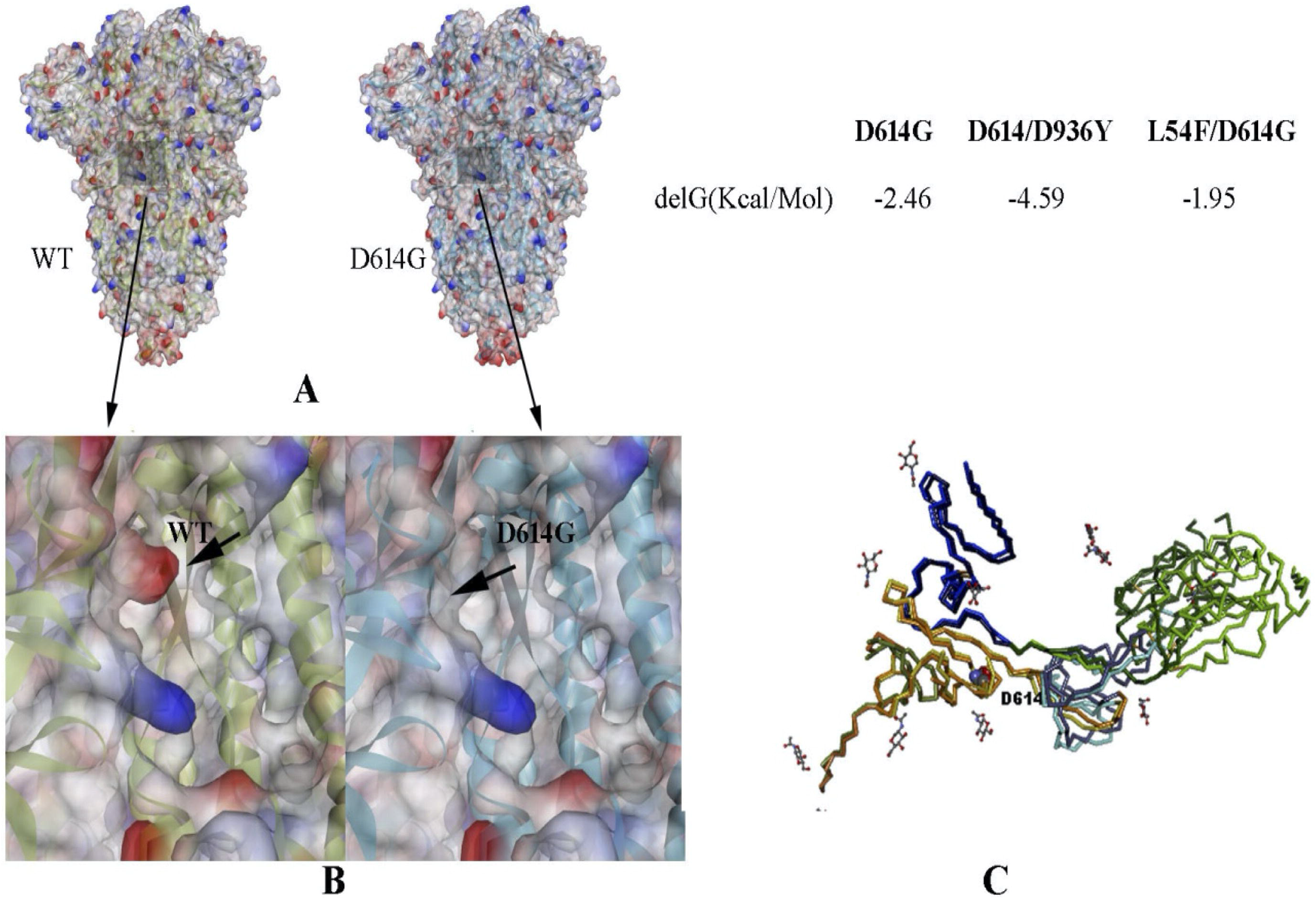
The implication of D614G mutation, **A.** Wild-type and D614G mutant structure represented as electro-static potential surface. The position of mutation is highlighted. **B.** A close view near the position of mutation. **C.** Superposition of a monomer of the DOWN/close wild-type (6VXX) and the UP/open (6VYB) of S protein showing a hinge-bending motion of RBD (green) around NTD linker (blue) and CTD2 (light blue). The location of D614 residue in CTD3 (orange) is indicated and represented as Van der Waal presentation. The neighbouring glycans are represented by stick model. Free energy change due to D614G single and D614G containing double mutants are given.

### Temporal and geographical distribution of wild type and mutant Spike glycoproteins

Among these multiple variants, the ones that are occurring in a large fraction of the samples can be said to have adapted, while those strains which only existed with very few samples were likely to get eliminated in the way of selective process and are not generally perceived among the emerging variants. This implies that the favourable variants should be associated with a greater stability and/or higher transmission rates of the SARS-CoV-2 proteins, while a decreased stability or transmission rate is expected in case of the minor variants.

Looking into the spatial and temporal distribution of these variants of S protein in Nextstrain and noting down the number of occurrences of each variant along with the country it originated with the corresponding date (Table 5), we find that the variant with D614G substitution is characterized by greater viability across different countries as seen over a span of time, first originating on 24^th^ December 2019 and prevailing since last recorded date. This mutation was also accompanied by L54F, D936Y and S939F in different isolates. However all these variants are observed in multiple samples, which show that the change at the 614^th^ residue is the impactful one which is imparting greater stability to the mutant protein. On the other hand the less stable mutants that were found only in a few samples did not show such prevalence, and were seen to be dwindling out with time.

**Table 5:** Geographical and temporal distribution of Spike protein mutants

| Mutations in Spike Protein |  | No. of Sample | COUNTRY | First Collection Date | Last Collection Date |
| --- | --- | --- | --- | --- | --- |
| A27V |  | 1 | USA | 23.03.2020 | NA |
| Y28N |  | 1 | Australia | 28.02.2020 | NA |
| T29I |  | 3 | Australia, USA, Netherlands | 21.03.2020 | 28.03.2020 |
| **H49Y** |  | **13** | **China, USA, Taiwan, Mexico, Australia** | **17.01.2020** | **26.03.2020** |
| **S50L** |  | **5** | **China, Singapore, Australia** | **28.02.2020** | **19.03.2020** |
| R408I |  | 1 | India | 27.01.2020 | NA |
| H519Q |  | 1 | Belgium | 29.02.2020 | NA |
| A520S |  | 2 | USA | 13.03.2020 | 03.04.2020 |
| A570V |  | 1 | China | 29.01.2020 | NA |
| **D614G** |  | **Innumerable** | **Across the globe** | **24.01.2020** | **Till date** |
| V772I |  | 1 | Turkey | 17.03.2020 | NA |
| F797C |  | 1 | Sweden | 07.02.2020 | NA |
| A930V |  | 1 | India | 31.01.2020 | NA |
| **D614G && L54F** | D/L | Innumerable | Across the globe | 24.12.2020 | 15.04.2020 |
|  | G/L | Innumerable | Across the globe | 24.01.2020 | 20.04.2020 |
|  | D/F | 1 | Netherlands | 31.03.2020 | NA |
|  | **G/F** | **5** | **France, USA, UK, Australia** | **07.03.2020** | **02.04.2020** |
| **D614G && D936Y** | D/D | Innumerable | Across the globe | 24.12.2019 | 15.04.2020 |
|  | G/D | Innumerable | Across the globe | 24.01.2020 | 20.04.2020 |
|  | **G/Y** | **11** | **Netherlands, Australia, Sweden, Wales** | **12.03.2020** | **02.04.2020** |
|  | D/H | 1 | Singapore | 05.01.2020 | NA |
| **D614G && S939F** | D/S | Innumerable | Across the globe | 24.12.2019 | 15.04.2020 |
|  | G/S | Innumerable | Across the globe | 24.01.2020 | 20.04.2020 |
|  | D/F | 1 | Switzerland | 26.02.2020 | NA |
|  | **G/F** | **7** | **France, Iceland, USA** | **04.03.2020** | **02.04.2020** |

\*Mutants with reduction in total free energies are presented in bold.

## DISCUSSION

We have performed a thorough mutational characterization encompassing the variations occurring in all ORFs of the SARS-CoV-2 genome. Among structural proteins, both the membrane and envelope proteins are more resilient to frequent mutations, while among non-structural proteins, ORFs 6, 7a and 10 shared similar behaviour to E and M proteins, with them being mostly conserved. This signifies that these proteins could have some essential functions, perhaps housekeeping roles that are critical to the virus, which is why these sequences cannot generally withstand any variations. In contrast, S, N, ORF3a, ORF8 and ORF1ab exhibits mutations. An intriguing feature for N protein that we noticed here was that both substitutions (R203K and G204R) were present simultaneously in 26 of the 28 samples, with only the 2 remaining samples lacking the G to R changes at the 204^th^ residue. Mutations in ORF8 showed a major substitution L84S and an accompanied substitution of V62L with few isolates with S24L substitution. Moreover, all of these changes are not independent with respect to one another, which is established from the fact that V62L is also accompanied by a corresponding substitutions S84L, with these mutations appearing to share a co-dependence with one another. We were fascinated to find that two major amino acid substitutions D75E and P971L of ORF1ab occurred in the same eighteen strains that harboured both of these mutations, and with no instance of any other strain having mutated at only one of these positions. These implicates that these two positions may have a linked relationship which could have been involved in some critical functions. Likewise, another clear-cut division of two variants was observed at the 4715^th^ position which possessed L and P variants. We can discern a possible link between this mutation and the one discussed at the 265^th^ position, both of which explicitly divided the isolates into two groups. Additionally we also detected two mutations at 5828^th^ (L to P) and 5865^th^ (C to Y) positions, and those strains that contained any one of these variants was also forced to accommodate the other variation, with no exception to this event being observed in any sample, which shows the complete linkage between the two changes. Combined analysis, only with the frequently mutated residues, identified at least 20 possible variants, among which 17 variants occurred at least more than one among the samples considered in this study. Frequent occurrences of some of the specific combination of mutations at 5 genes clearly indicated their direct or indirect interaction leading to stability, adoptability, viability and transmission efficiency of the virus. Less frequently occurred variants might have eventually lost due to their low transmission efficiency or less adoptability in nature. Country specific under- and/or over-sampling could be a confounding factor for this variation. However, our observation with ∼660 samples showed excellent concordance with the data generated from ∼10,000 samples, suggesting the generality of this observation.

The contribution of mutations to the stability and function of the gene product, which depends on its location, including interaction with other viral or host molecules, may determine the viability of the mutant. Absence of any charge reversal, either among SARS-COV-2 mutants or SARS-COV or bat and pangolin corona viruses, and low frequency of mutations with change in charge state underscore that charge state plays an important role in viability of variants. The high viability of D614G mutant of S-protein seemed to be attributable to miniscule changes in partitioning energy as well as the fact exposed aspartate side chain located on a flexible loop in a relatively hydrophobic environment was not involved in any H-boding while it's substitution by glycine could facilitate the movement of the hinge. Compensatory effects of additional H-bond could be a plausible explanation for relatively high frequency of L84S (ORF8) and Q57H (ORF3) mutants. However, these simple parameters could not explain low abundance of V483A (S-protein), V62L (ORF8), V5550L and D75E (ORF1ab), R203K (N) or discrepancies in abundance of L3606F vs F6158L and P/L mutants (at 4715 and 5528 vs 971 position) of Orf1ab. Unlike D614A, V483A mutant is a part of the crucial receptor binding domain. The tighter binding (4-10-fold compared to SARS-COV-1) of the S1-CTD to hACE2 receptor [22, 37] has been attributed to the enhanced infectiousness of SARS-COV-2. Thus, low frequencies of the V483A as well as other S1-CTD mutants seem, primarily, attributable to their role in interaction with the host receptor. It is possible that V62 (ORF8); R203 (N); V5550, D75, P971, L3606 and F6158 (ORF1ab) positions are also associated with crucial functional roles beyond stability.

Two variants with co-occurring mutants, were more prevalent than the wild type variant. The most prevalent variant showed co-occurrence of P5828L and Y5865C in ORF1ab. The ability of proline to introduce kink in the structure - often in turns and loops close to surfaces and the tendency of the upstream cysteine to be modified if exposed or form S-S bond if buried may explain the co-occurrence. The next prevalent variant showed co-occurrence of T265I and P4715L in ORF1ab. This might be indicative of these hydrophobic substitutions coming closer in the tertiary structure and stabilizing it through van der Waals interaction. The co-occurrence of these ORF1ab mutants with D614G (S-protein) and Q57H (ORF3a) is suggestive of functional interaction among these proteins. However, these interpretations are contingent upon the reported mutation frequencies being representative of the actual variant distribution and certainly begs more investigation and analysis.

Among the structural proteins which were mostly conserved, only the spike protein was found to be associated with several mutations including a dominant mutational variant at the 614^th^ position. We have investigated the thermodynamic stability of the variants to establish those variants which are correlated with greater stability and sustainability. Those strains that corresponded to structures with low stabilities were consequently found to have low transmission capabilities as verified in the Nextstrain data. We have identified several mutants with stable structures, including mutations at positions 49, 50, 54, 614 and 936 and have verified that these variants are enduring among the general population over time, with D614G be the most viable among them. Although, some of the mutated residues of spike protein showed greater reduction of total free energy compared to D614G substitution, but their spatio-temporal distribution and number of isolates are comparatively lower than the substitution at 614. It clearly suggests that spike protein alone is not the determining factor of the stability, adaptability and transmission efficiency of the virus. Specific combination of all frequently mutated variants might be necessary for the prediction of viability of the viral variants. However, considering only the disparity in the effectiveness of transmission among the different spike protein variants, we have two important suggestions to the different nations in tackling and curbing the spread of COVID-19 with greater efficacy. First and foremost, the mutational profile of a patient found to be COVID-19 positive needs to be analysed, specifically at these key sites of five proteins, either by Sanger sequencing or designing probes corresponding to these regions. Thereafter, a model can be predicted using the patients severity and transmission of infection among the contacts for each combination of frequently mutated residues. Though one could argue that as the sequencing of the viral genome had been carried out at different time-points in different countries, with some countries like China imposing higher levels of quarantine measures at an earlier time compared to other countries [38], our interpretations may not have 100% accuracy. However, our hypothesis and interpretation of the mutations show good concordance as evidenced from the Nextstrain data. Further research on identification of mutational status SARS-CoV-2 infected individual and determination of infection among their contacts might help to substantiate the idea of correlation between genotypes with survivability and transmission of different strains. In conclusion, we maintain the belief that the propositions voiced here if followed adequately can work to curb the spread of the disease with much higher success.

## Acknowledgments

We would like to acknowledge Prof. Nitai P. Bhattacharya, retired professor, SINP Kolkata for his critical comments and evaluation of the manuscript.

